# Hybrid *Mimulus* flowers attract a new pollinator

**DOI:** 10.1101/2023.09.19.558487

**Authors:** Foen Peng, Xiaohe Sun, Claudia van Vloten, Jude Correll, Marlena Langdon, Weerin Ngochanthra, Karl Johnson, Suzanne Amador Kane

## Abstract

Hybridization is common in flowering plants and is believed to be an important force driving adaptation and speciation. The hybrid flowers often have new trait combinations, which, theoretically, could attract new pollinators. In this study, we found that the hybrids between a hummingbird-pollinated species *Mimulus cardinalis* and a self-pollinated species *Mimulus parishii* attract bumblebees (*Bombus impatiens*), a pollinator not attracted to either of the progenitor species. This novel attraction is explained by several floral traits in hybrids, including petal color, nectar concentration, and corolla size. We also found that the *YUP* locus, previously shown to be important in bee attraction in other *Mimulus* species, also played an important role in this novel attraction. Hybrid plants’ attraction to new pollinators could be an underexplored avenue for pollinator shift and speciation.

## Introduction

Hybridization occurs naturally in many taxa. When species hybridize, two diverged genomes come together in the same nucleus, which brings both challenges and opportunities. For example, the diverged chromosomes often cannot be separated properly in meiosis due to poor homology, which leads to low quality gametes and reduced fertility. Also, DNA methylation may not be regulated properly in the hybrid genome, resulting in aberrant activity of transposable elements and high mutation rates (McClintock 1984). However, in some cases, the hybrids could have some unique advantages. For example, hybrids could benefit from the genetic materials introgressed from a different genome to deal with challenges in the environment (Meier et al. 2017; Grant and Grant 2019; Pardo-Diaz et al. 2012). Also, hybrids often have novel phenotypes which could be advantageous in newly opened ecological niches inaccessible to either parent (Selz and Seehausen 2019).

In flowering plants, hybridization is common and widespread (Whitney et al. 2010). Hybrid plants often show a combination of traits from both parents, which is particularly interesting from the perspective of pollinator attraction, as it is unclear which groups of pollinators the hybrids would attract. Many plant species specialize to attract a particular group of pollinators, *e*.*g*. hummingbirds or bumblebees, to maximize pollination efficiency. These plant species often evolve a suite of floral traits, involving color, shape, scent *etc*., described as pollinator syndromes, to meet the needs of the pollinator for which they are specialized (Fenster et al. 2004). For example, hummingbird-pollinated species often have flowers that are red, with a long corolla tube or nectar spur to fit the bird’s beak, and produce large volumes of diluted nectar. In contrast, bumblebee-pollinated flowers often have blue or pinkish color, a wide corolla to provide a platform for bees to land, and small amounts of concentrated nectar. When two species with different pollinator syndromes hybridize, the resulted hybrids are expected to have a combination of traits from the two syndromes.

There are several possible outcomes of such hybridization in terms of pollinator attraction (Grant 1993). First, the hybrids may not attract either pollinator that the progenitors specialized for, and hence would not persist. Second, if the hybrids are still visited by pollinators attracted by one of the progenitors, they could merge into that progenitor species through backcrossing. In this process, hybrid introgression could occur, meaning the hybrids bring genetic materials from another species into the backcrossed progenitor species. Third, the hybrids could persist as a hybrid swarm if both progenitors’ pollinators continue visiting the hybrids. Finally, the hybrids could attract a new pollinator that neither progenitor attracts. The latter is possible because the hybrids have a variety of novel trait combinations through chromosome assortment and recombination, as well as some transgressive traits that neither parent has (Rieseberg, Archer, and Wayne 1999). These various new traits and new trait combinations could be explored by other pollinators in the community (Vereecken, Cozzolino, and Schiestl 2010). For the hybrids, being pollinated by a new pollinator could be advantageous, as the hybrids would avoid direct competition for pollination service with their progenitors. In addition, being pollinated by a new pollinator also means the gene flow between hybrids and progenitors become limited. Therefore, pollinator-mediated reproductive isolation could lead to the formation of a new hybrid species, a case of homoploid speciation (Ungerer et al. 1998).

The genus *Mimulus*, including over 150 species, is well known for its abundant floral variation and frequent pollinator syndrome transitions (Wu et al. 2008). Closely related *Mimulus* species are often interfertile. Recent work suggested that hybridizations have occurred frequently among closely related *Mimulus* species in the wild (Nelson et al. 2021). This raises the question of whether hybridization facilitates pollinator syndrome transitions. In this study, we set out to test this possibility. We focused on two sister species (Fig 1): a) *Mimulus cardinalis*, which is specialized for hummingbird pollination, has red and unscented flowers with abundant diluted nectar; and b) *Mimulus parishii*, which is self-pollinated and has miniature-sized pink flowers with no scent or nectar. Both species are distributed in the west coast of the United States and their ranges overlap in southern California, where existing hybrids have been reported (https://www.inaturalist.org/observations/91539400). In greenhouse conditions, the two species produce vigorous and fertile hybrids.

**Figure 1.**
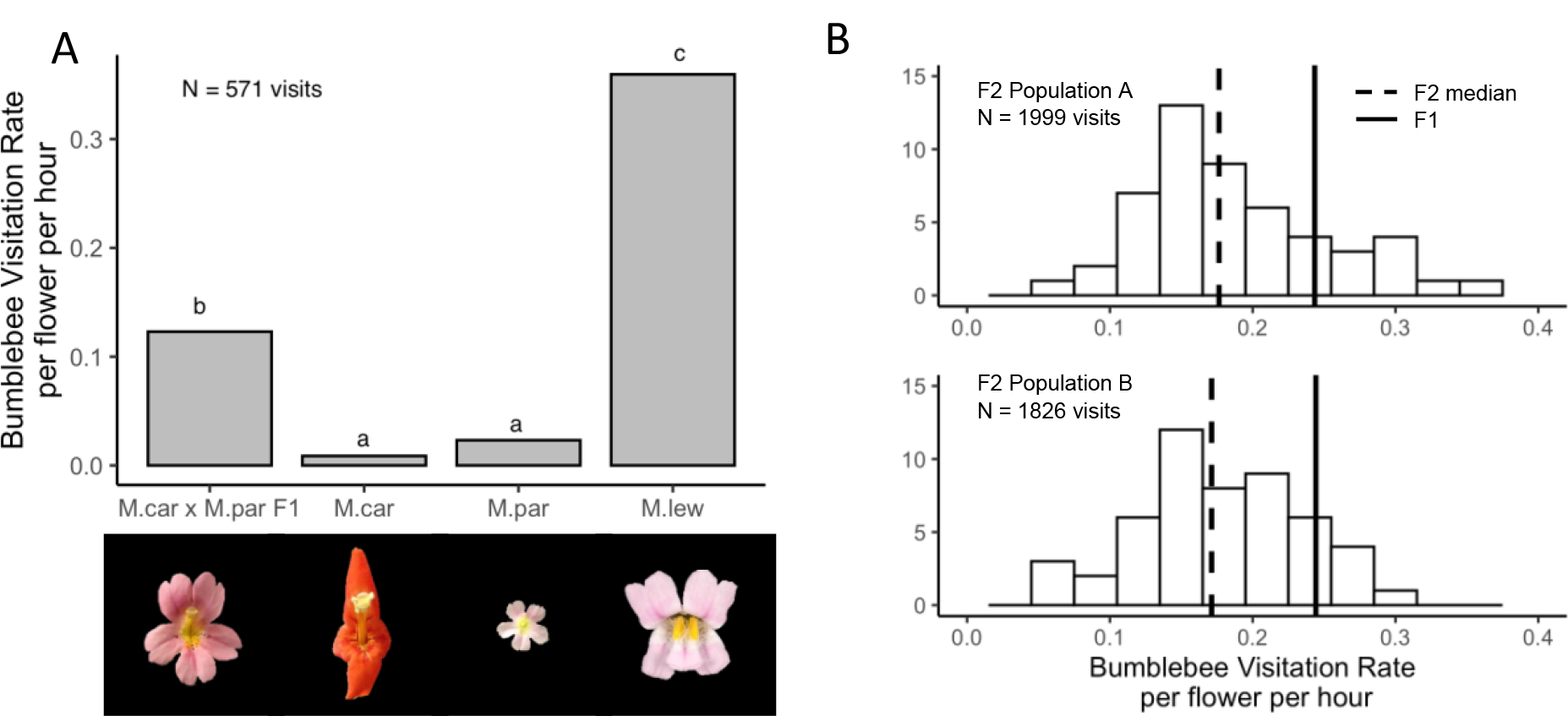
A. Bumblebee visitation rates in Pollination Experiment 1. The images of the respective flowers are below the bar graph. In the wild, *M. cardinalis* is hummingbird-pollinated, *M. parishii* is self-pollinated, and *M. lewisii* is bumblebee-pollinated. B. Histogram of bumblebee visitation rates in Pollination Experiment 2. The vertical axis represents the number of individuals in each bin of visitation rate. The dashed and solid vertical lines mark the median visitation rates of the F2 and the visitation rate of F1s, respectively.

This study was motivated by field observations by one of the co-authors (Peng). He observed that the F1 hybrids between *M. cardinalis* and *M. parishii* were frequently visited by wild bumblebees when the plants were put outside of the greenhouse for an exhibit in Washington state in the U.S., where neither parental species exists naturally. In contrast, none of the parental species next to the F1s received visits from the bees. This observation suggests that the F1s might contain a trait combination that is favorable for bumblebee pollination, despite neither parent attracting bees.

In this study, we tested two hypotheses. First, we hypothesized that the F1 hybrids’ attraction to bumblebees was not accidental, instead, it was because of bumblebees’ preferences to the F1s’ new floral trait combination. To test this hypothesis, we conducted a pollinator choice assay with *M. cardinalis, M. parishii*, and their F1 hybrids, using commercially purchased bumblebees (*Bombus impatiens*). Second, given that pollination is a complex process, involving visual attraction, the fit between the animal body and the corolla shape, appropriate rewards, *etc*., we hypothesized that there were multiple floral traits in the hybrids contributing to this novel attraction. To test the second hypothesis, we conducted another pollinator choice assay using F2 hybrids, as F2 hybrids segregate a variety of floral traits while the F1s are phenotypically uniform.

## Results

### 1. F1 hybrid plants were very attractive to new bumblebee pollinators

We first performed a greenhouse pollination experiment (Pollination Experiment 1) with commercially purchased bumblebees (*Bombus impatiens*), including an equal number of plants from *M. cardinalis, M. parishii*, the F1 hybrids between these two species, and the bumblebee specialist *M. lewisii* as a positive control. Results from this experiment showed that the F1 hybrids were much more attractive to bumblebees than either parent (Fig. 1A). A total of 571 visits were recorded in 6.5 total observation hours spanning 5 experiment days. F1s were significantly more attractive to bumblebees than either *M. cardinalis* (p<0.001, Chi-squared test, p value adjusted for multiple comparisons) or *M. parishii* (p<0.001, Chi-squared test), but less attractive than the bumblebee specialist *M. lewisii* (p<0.001, Chi-squared test) (Fig. 1A). *M. cardinalis* and *M. parishii* were similarly unattractive to bees (p=0.29, Chi-squared test).

### 2. Several floral traits correlated with bee visits, with petal color being the most significant factor

We then performed Pollination Experiment 2 in order to infer the floral traits correlated with bee attraction. In this experiment we studied two replicate groups, A and B, of F2 hybrids between *M. cardinalis* and *M. parishii*, which segregated in a variety of floral traits. Each replicate group had 50 F2s, and 2 F1s were included as internal controls as their phenotypes were uniform. We recorded 1999 visits for group A (a total of 12.0 observation hours over 4 experiment days) and 1826 visits for group B (a total of 10.3 observation hours over 4 experiment days). The two groups showed similar distributions of visitation rates (two-sample Kolmogorov-Smirnov test, p = 0.97. Fig. 1B). While a majority of F2s were less attractive to bees, compared to F1s, a few F2s were more attractive. Specifically, the F1s corresponded to the 82th percentile and 86th percentile in terms of attractiveness in all the plants in group A and group B, respectively.

We measured many floral traits for each of the F2 plants and performed a negative-binomial regression to analyze their correlation with bee visits. We found that petal color (measured as the average R, G, B values of a flower’s digital photograph in the front view) had the most significant impact on bumblebee visits, followed by nectar concentration and projected area. Flowers with more blue hue (z value = 6.05, p<0.001), higher sugar concentration (z value = 2.44, p=0.01) and larger overall projected area (z value = 2.23, p=0.03) were visited more frequently by bees. None of the other traits, such as pistil length, nectar volume, petal reflex, and petal spot sizes, had a significant correlation with bee visitation.

Consistent with this result, we found that F1 flowers have a combination of high blue hue (26.8% in F1 vs. 11.2% in *M. cardinalis* and 29.8% in *M. parishii;* the percentage was calculated as the B value divided by the sum of R, G, and B values), high sugar concentration (27.9% in F1 vs. 13.9% in *M. cardinalis* and 1.3% in *M. parishii*), and large projected areas (4.6 cm^2^ in F1 vs. 3.38 cm^2^ in *M. cardinalis* and 1.14 cm^2^ in *M. parishii*). This is consistent with the relatively high attractiveness of the F1s in both Pollination Experiment 1 and Pollination Experiment 2.

### 3. The color contrasts in the F2s were conspicuous enough for bees to detect and the most-visited F2s appeared brighter

Given the importance of petal color for bee pollination preferences, we modeled the appearance of flowers from the perspective of bee vision for the 10 most visited and 10 least visited F2 individuals in Pollination Experiment 2. These two groups differed in their reflectance spectra (Fig. 2A). Notably, the most visited F2 petals had stronger reflection in the 400-500 nm region, which overlapped with the sensitivity peak of bumblebee blue photoreceptor (424.1nm) (Skorupski and Chittka 2010). We analyzed the quantitative differences between the two flower groups using two commonly used vision models: the Color Hexagon model (Chittka 1992) and the Receptor Noise model (Vorobyev et al. 2001) (Fig. 2C). Based on the Color Hexagon model (Fig. 2B), we calculated the green contrast (*i*.*e*. the perceptual contrast between two colors detected by the green photoreceptor adjusted to the background) and brightness differences (*i*.*e*. the sum of the three photoreceptors excitations) between the most- and least-visited F2s. We found that the most visited F2s had stronger green contrast (t-test, p=0.049) and would appear brighter to bees (t-test, p=0.007). The color contrast analysis based on Receptor Noise model indicated bees can distinguish the most- and least-visited F2 flowers from each other. The color difference between the most- and least-visited groups was 4.1 JND (Just Noticeable Difference), which was greater than the 1 JND detectability threshold. Both the most- and least-visited flowers could be distinguished from the green foliage background by bees (Fig. 2C). The flower images generated by multispectral imaging (Fig. 2D) showed that the most- and least-visited F2 flowers appeared in different colors in bee vision and the most visited ones were brighter than the least visited ones, which was consistent with the quantitative analysis based on reflectance spectra. Neither the progenitor species nor the hybrids had UV reflection in their flowers.

**Figure 2.**
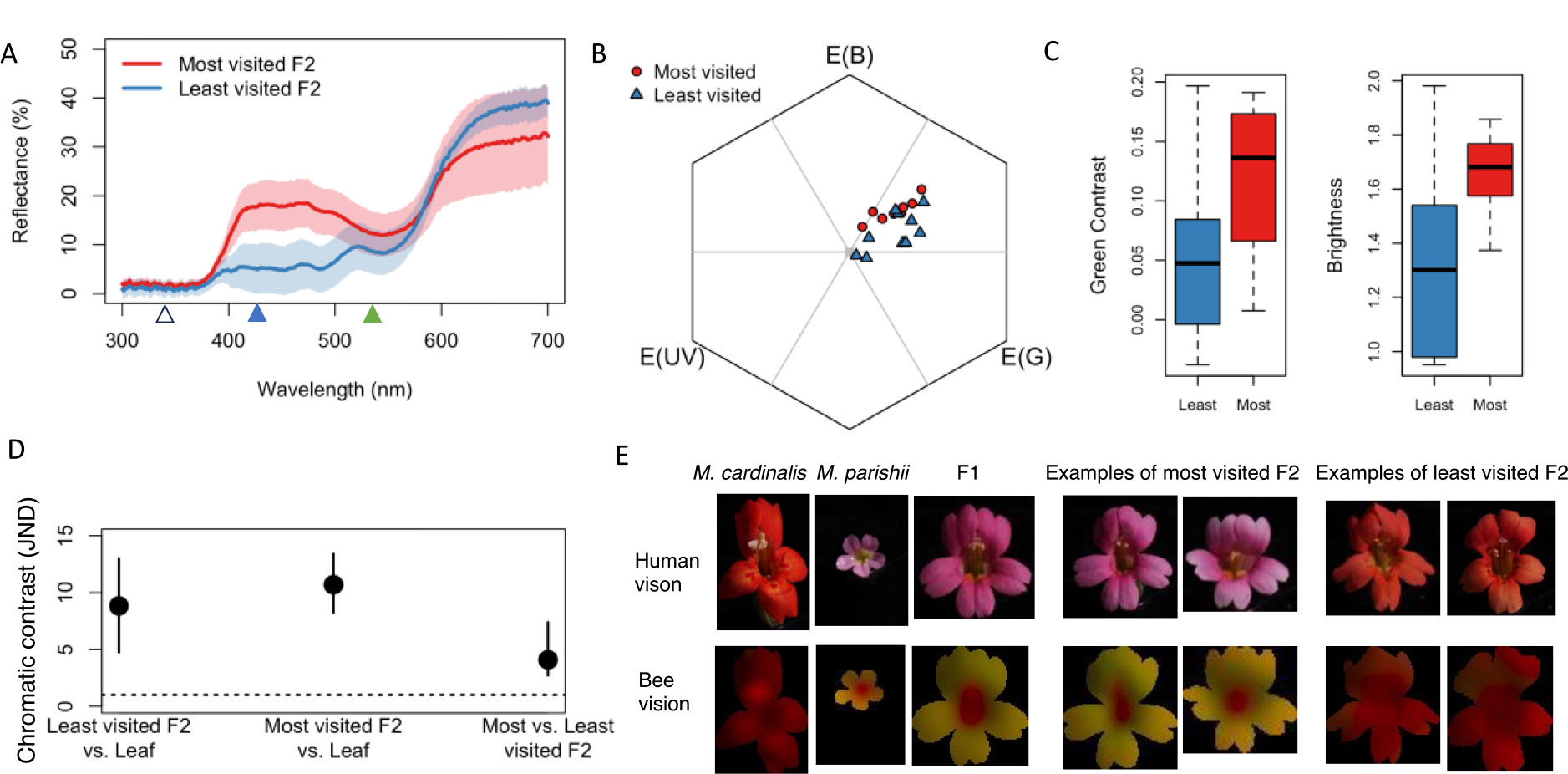
A. Reflectance spectra of the most and least visited F2 hybrids in Pollination Experiment 2. The colored triangles on the horizontal axis mark the peak sensitivity of the three photoreceptors (left to right: UV, Blue, and Green) the bumblebee species *Bombus impatiens* used in this study. B. Petal color projection using the Color Hexagon model. C. Green contrast and brightness differences between the most and least visited F2s, calculated using Color Hexagon model. D. Chromatic contrast among the most and least visited F2s and leaves, calculated using the Receptor Noise model. D. Representative flower images shown in both human vision and false color images for the bee vision model taking into account blurring due to visual acuity.

### 4. *M. parishii YUP* allele is favored by bumblebees

Previous work showed that the *YUP* supergene on chromosome 4 could influence petal color and pollinator preference (Liang et al. 2023; Bradshaw and Schemske 2003). This genomic region includes several genes involved in petal color pigmentation regulation, such as *PELAN* and *YUP* (Liang et al. 2023). Through AFLP genotyping, we found that the most visited F2s were more likely to have at least one allele inherited from *M. parishii* parent (9 out of 10 were P_ genotype. P indicates *M. parishii YUP* allele while C indicates *M. cardinalis YUP* allele), while the least visited F2s were more likely to be homozygous *M. cardinalis* genotype (8 out of 10 were CC genotype).

We then extended our genotyping efforts to all the F2s and analyzed the impact of *YUP* on bee visitation. The F2 plants with at least one *M. parishii YUP* allele (PC or PP genotypes) were visited more frequently by bees (Kruskal-Wallis rank sum test, p<0.001. Dunn’s *post hoc* test, PP-CC p=0.008, PC-CC, p<0.001), and there is no significant difference between PP and PC genotypes (Dunn’s post hoc test, PP-PC p=0.67) (Fig. 3). As expected, we found that *YUP* locus significantly impacted petal color (Kruskal-Wallis rank sum test, relative red p<0.001 relative blue, p<0.001), with CC genotype having smaller B values and higher R values, compared to PC or PP genotypes (Fig. S1). But in addition, this genomic region also significantly influenced a range of other floral traits, such as petal shape (*e*.*g*., corolla tube length) and nectar volume (Fig. S2).

**Figure 3.**
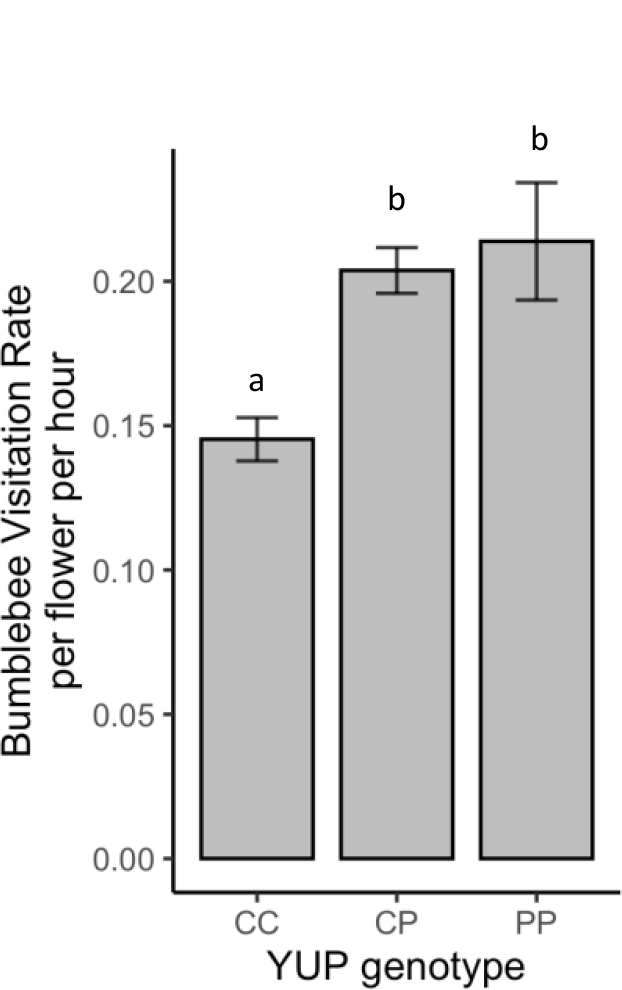
Impacts of F2 *YUP* genotype on bumblebee visitation in Experiment 2. CC: homozygous *M. cardinalis* genotype; PP: homozygous *M. parishii* genotype; PC : heterozygous.

## Discussion

Hybrids with novel trait combinations are presumed to be able to explore novel ecological niches. In the case of pollination, this means hybrids with new floral trait combinations may attract new pollinators and promote reproductive isolation and speciation, but existing case studies supporting this are scarce (Rezende et al. 2020). Here we showed that the hybrids between hummingbird-pollinated *M. cardinalis* and self-pollinated *M. parishii*, are attractive to new bumblebee pollinators. The floral traits in the hybrids important for bee attraction comprise high nectar sugar content, large landing area, and most importantly, enhanced blue hue. We also found that the *YUP* supergene (Liang et al. 2023; Bradshaw and Schemske 2003), previously known to impact petal color and pollinator preference in a different species pair (between *M. cardinalis* and *M. lewisii* (Schemske and Bradshaw 1999)) also significantly impacts bee visitation in the offspring of the cross between *M. cardinalis* and *M. parishii*.

As expected, the hummingbird-pollinated *M. cardinalis* is not attractive to bumblebee pollinators. Its nectar is hard to access for bumblebees due to the narrow corolla opening, despite its large overall flower size. Also, the low sugar concentration of its nectar may not be motivating for bees even if they could access it. In addition, its scarlet petal coloration appears relatively dark in bee vision which may not help bees efficiently locate flowers. The self-pollinated *M. parishii* is also not attractive to bumblebee pollinators for different reasons. Its flower is too small to provide a stable landing platform for bumblebees and it has little to no nectar to reward a pollinator. In contrast to both parents, the F1 hybrid is much more attractive to bumblebees. It has a combination of large corolla size, a trait likely inherited from *M. cardinalis*, magenta petal coloration, a trait likely inherited from *M. parishii*, and high nectar sugar concentration, a transgressive trait that neither parent had. Particularly, the magenta coloration turned out to be important as it reflects more blue light around the 400-500nm region where the blue photoreceptor of bees detects, making the flower much more visible to bees in a green foliage environment. Overall, the F1 hybrid between *M. cardinalis* and *M. parishii*, to some extent, mimics the bumblebee-pollinated *M. lewisii* (*e*.*g*. both the F1 and *M. lewisii* have magenta unfolded petals and a small volume of highly concentrated nectar), which is not surprising given the recent divergence history of these species (Beardsley, Yen, and Olmstead 2003; Nelson et al. 2021). However, the F1 hybrids traits are still not optimally suited for bumblebee pollination, as *M. lewisii* is about 3 times more attractive to bumblebees than the F1. The experiment with F2 individuals also showed that as phenotypic traits begin to segregate, while the majority of individuals were less attractive to bumblebees than the original F1 hybrid, some plants have higher visitation rates. It would be interesting to know what the outcome would be if the hybrids were under consistent bumblebee selection over many generations, for example, whether it would evolve towards a more *M. lewisii*-like floral appearance.

The new combination of floral traits as a result of hybridization is key to this novel attraction. Hybrids, carrying a mixed genome and new combinations of floral traits, might have an advantage when a new ecological niche becomes available (Selz and Seehausen 2019; Feller et al. 2020). Many works have shown that hybridization could fuel speciation and adaptive radiation, such as in cichlid fish (Meier et al. 2017), Darwin’s finches (Grant and Grant 2019), and *Heliconius* butterflies (Pardo-Diaz et al. 2012). In flowering plants, hybridization is particularly common and suggested to play a significant role in plant speciation (Soltis and Soltis 2009). Pollinator preference is an important form of pre-zygotic isolation in plant speciation. However, there is little knowledge about how hybridization influences the frequently observed pollinator shifts between closely related species (Rezende et al. 2020). In our study, we observed a stark contrast of attractiveness to bumblebees between the hybrid and its progenitor species, which suggests the possibility of pollinator shift. It is reasonable to expect that if it occurs in nature, the pollinator-mediated reproductive isolation could significantly reduce gene flow between the newly formed hybrids and parents, thus promoting new species formation.

Flower adaptation to animal pollinators is a good example of multi-trait adaptation, as it involves many floral traits, such as color, scent, and shape, evolving coherently. The genomic basis that enables this type of complex adaptation is interesting but hard to study due to the involvement of many genes. In this study, we tested the effect of one locus, *YUP*, that’s previously known to impact pollinator preferences (Schemske and Bradshaw 1999) and found that *YUP* also had a significant impact on bumblebee pollination in our experiment. The *YUP* region includes many genes controlling flower traits. These genes form a tightly-linked genomic island, which was hypothesized to be advantageous in polygenic adaptation, as the suitable alleles are always inherited together as a unit (Schwander, Libbrecht, and Keller 2014). Some of the genes in *YUP* region have been identified, such as *YUP* gene, a siRNA gene repressing carotenoid pigment synthesis, and *PELAN* gene, encoding a MYB transcription factor controlling anthocyanin pigment production in the petal lobe area, while the others remain unknown, such as the genes affecting petal shape and nectar volume. Given the considerable impact of petal color on bee visits and the strong effect of *YUP* gene on petal color (a binary determinant of petal carotenoid production), we infer that the *YUP* gene may be the most important gene in this region that influences bee attraction in our experiment. While the *YUP* region was presumed to be under strong directional selection between *M. lewisii* and *M. cardinalis* to maintain the haplotype structure (Bradshaw and Schemske 2003), there would not have been similar selection pressure between *M. cardinalis* and *M. parishii*, as neither of the species are bee-pollinated. The reason why *M. parishii* has this bee-attracting *YUP* allele could either be a result of incomplete lineage sorting (ILS) from the shared ancestor with *M. lewisii*, or that it was introgressed from *M. lewisii* after a historical hybridization event. While we have confirmed the role of *YUP*, further efforts will be needed to characterize other genes which also contributed to this novel attraction, such as the genes influencing nectar concentration and petal projected area.

In this study, we witnessed an unexpected relationship between hybrids and a new group of pollinators. Subsequent experiments revealed the mechanisms underlying this novel attraction, including hybrids’ novel trait combination and transgressive traits at the phenotypic level, as well as supergene and the inheritance of advantageous haplotype as a result of ILS or introgression at the genotypic level. While the observation that hybrids attract a new pollinator was initially surprising to us and relatively uncommon, the underlying mechanisms are all well-known and widely documented (Rieseberg, Archer, and Wayne 1999; Bell and Travis 2005; Feng et al. 2022; Rivas-González et al. 2023; Suarez-Gonzalez, Lexer, and Cronk 2018; Hedrick 2013). Therefore, we surmise that similar phenomena in other taxa are waiting to be discovered. Eventually, with further field work in natural habitats, it will be interesting to know how likely the novel pollinator attraction could result in pollinator shift, and how common this is as an avenue leading to homoploid speciation.

## Materials and methods

### Plant growth

The F1 hybrids were generated by a cross between the *M. cardinalis* inbred line CE10 (pollen donor) and a *M. parishii* inbred line (pollen receiver). The F1s were self-pollinated to generate the F2 seeds. *M. lewisii* LF10 inbred line was also planted for pollination experiments. All the plants grew in the Haverford College, PA greenhouse in Spring 2022 under long day condition (16hr light / 8hr dark). The plants were watered daily with fertilized water.

### Floral traits measurement

Newly opened flowers were labeled every day during the measurement period and only flowers that were 1-2 days old were used in trait measurements. A total of 15 traits in 5 plants of each parental species, 5 F1s, and 100 F2 plants were measured, including corolla tube length (CTL), corolla tube width (CTW), corolla lobe length (CLL), corolla lobe width (CLW), long stamen length, pistil length, stalk length, nectar volume, nectar concentration, projected area, petal color (R, G, B), petal reflex and nectar guide spot size. Flower volatiles were not collected because neither parent species (Peng, Byers, and Bradshaw Jr. 2017) or the F1s (unpublished data) produce bee attracting volatiles. We used a digital caliper for all the shape measurements except projected area, which was estimated from images. We measured two flowers per plant for shape measurements. For example, corolla tube length, CTL, was measured as the distance from the receptacle to the petal split between dorsal and lateral petals. For a detailed explanation of all the shape variables, see (Chen 2022). Nectar volume and nectar concentration were also measured in two flowers per plant. Nectar volume was measured with 25uL Microcaps disposable glass micropipettes and nectar sugar concentration was measured with an Eclipse low volume refractometer. Nectar from flowers with a volume less than 2 uL were diluted with 2uL distilled water before measurement. Then, the before-dilution concentration for those flowers was calculated. The projected area and RGB color were measured on a front view photo of one flower for each plant. We took the flower photos with an Olympus TG-6 camera in a dark room with constant lighting and made sure that the photos were not overexposed. We used the tracing tool in ImageJ to find the outline of the flower then used the ROI tool to calculate the area and the average R, G, and B color values inside the flower region. Although digital photograph taken by commercial camera may distort color information, recent work showed that they still capture a significant amount of color information and could generate comparable results with traditional spectrometry approach (Laitly et al. 2021). Finally, the petal reflex was visually scored on 3-5 flowers per plants into three categories: complete reflex, intermediate reflex, or no reflex. Similarly, nectar guide spot size was visually scored to large connected areas, intermediate, or small unconnected dots.

### Pollination experiment setup

Two bumblebee hives (*Bombus impatiens* Natupol Excel Startup) were purchased from Koppert in Spring 2022. The hives included a queen and over 100 workers. The hives had access to sugar water and were fed with fresh pollen (also from Koppert) every other day. The hives were acclimated to a flight cage (Growneer crop cage ordered from Amazon), which was made of netting materials with a dimension of 3m (L) x 2m (W) x 2m (H), for a week before the pollination experiment. The flight cage was put outside of the Haverford College greenhouse so that the bees foraged on flowers exposed to natural sunlight. During acclimation of the bees, 30-40 F2 plants each day were placed in the cage, none of which were used in the actual experiment later. Sugar water supply to the hives was cut off for 16 hours before each experiment day to motivate the bees to search for food. During the experiment, the bees in the hive were allowed to freely forage within the flight cage and return to the hive. The plants in the cage were placed in a 6 x 9 array and the positions were randomized at the beginning of every experiment day (Fig. S3). The number of flowers on each plant was counted before experiments started. The experiments were conducted in late April to early May 2022. Each experiment day lasted 4-6 hours, depending on how active the bumblebees were. Typically, there were 5-15 bees foraging in the cage at any given point. The experiment was observed by two observers for multiple 20-min windows between 9am and 4pm. During those periods, the observers record the number of visits to each plant. A visit is counted when the bees made physical contacts with anther or stigma, or the bees crawled into the corolla tube to access nectar.

### Pollination Experiment 1

A total of 12 plants, including 3 *M. cardinalis, 3 M. parishii, 3 of* the F1 hybrid between *M. cardinalis* and *M. parishii*, and 3 *M. lewisii*, were used in each day’s experiment. The Pollination Experiment 1 include 5 days of observations.

### Pollination Experiment 2

A total of 104 plants were used for Pollination Experiment 2, including 100 F2 plants and 4 F1 plants. We randomly divided the plants into two groups: A and B. Each group had 50 F2 and 2 F1 plants and was exposed to bees for 4 experiment days.

### Bee vision modeling

We took reflectance spectra for the 10 most visited and 10 least visited F2 flowers in Pollination Experiment 2. This sample size was sufficient to represent the two ends of the visitation spectrum for quantitative color analysis (Dalrymple et al. 2015). The petal lobe area of flower samples was illuminated using a PX-2 pulsed Xenon light source, triggered by a 200Hz 5V TTL function generator pulse. An Ocean Optics USB2000+RAD spectrophotometer was used for data collection with OceanView software parameters: reflectance mode, 30ms integration time, averaging 5 consecutive scans, and a pixel boxcar average of 3. Reflectance spectra were processed using the R packages lightr and pavo (Maia et al. 2013). We used 347, 424 and 539 nm for UV, blue and green receptors to model the color hexagon model (Skorupski and Chittka 2010). For the color hexagon model, we followed (Martins et al. 2021) to calculate the green contrast and brightness: the green contrast was calculated by subtracting 0.5 from the green photoreceptor excitation and the brightness was calculated as the sum of the three photoreceptors excitations. For the receptor noise model, we used *Apis mellifera* noise parameters, with the receptor density ratio 0.85:4:1 and the weber fraction of 0.12 (Vorobyev et al. 2001). Garcia *et al*, (Garcia, Shrestha, and Dyer 2018) suggested that these noise parameters better represent behavioral data for bumblebees than the receptor-noise model constructed for bumblebees. We used a Canon EOS 7D camera for multispectral imaging, which had been full spectrum converted by removing its internal infrared-UV filter. We used an El-Nikkor 80mm/5.6 lens. One image was taken with a Baader visible-pass filter, and another with a Baader UV-pass filter. The flower samples were illuminated by an Iwasaki EYE ColorArc bulb. The MICA package in ImageJ was used for multispectral image processing (Caves and Johnsen 2018; van den Berg et al. 2020; Troscianko and Stevens 2015).

### Genotyping

A pair of primers was designed to perform the Amplification Fragment Length Polymorphism (AFLP) genotyping for the *YUP* locus. (Forward: TGGGTTGGTTATGGTCGATT; Reverse: CGTGAAAGCTTTACACATGTCT). It amplified a region close to YUP siRNA gene sequence (∼2kb upstream). The amplicon from *M. parishii* allele was 312 bp long while the amplicon from *M. cardinalis* allele was 470 bp long. We used GoTaq Master mix (Promega) for the PCR reactions. The following PCR protocol was used: step 1, 95°C for 3 min; step 2, 95°C for 30 s; step 3, 55°C for 30 sec; step 4, 72°C for 30 sec; 40 additional cycles of steps 2 to 4; finish at 72°C for 5 min. The PCR product was stained with EtBr and run on a 1.4% agarose gel to visualize.

### Data analysis

Data analysis was conducted with R 4.3.1 (R Core Team 2021). In order to calculate the bumblebee visitation rate, which was defined as visits per flower per hour, we divided the total number of visits over all experiment days by the total number of flowers and the total number of observation hours. For Pollination Experiment 1, Chi-squared test was used to evaluate the difference in attractiveness between different groups. The p value was adjusted using Bonferroni method to account for multiple comparisons. For Pollination Experiment 2, we tested whether the distribution of visitation rate between the two groups were similar with two-sample Kolmogorov-Smirnov test. Then we used a Mixed-Effect Negative Binomial Model to analyze the impacts of floral traits on the number of visits each F2 plant received, with the group assignments and the experiment dates as nested random effects variables. (Formula: visits∼ No_total_flowers + pst_len + stk_len + nec_vol + nec_con + CTL + prj_area + B + G + dot_size +(1|Group/Date)). We used two-sample T test to test whether there were significant differences in green contrast and brightness between the most and least visited F2s. For the genotyping results analysis, we used Kruskal-Wallis rank sum test and Dunn’s post hoc test to test whether there are significant differences in color value or visitation among the different genotypes.

## Acknowledgement

We would like to acknowledge all the undergraduate students in Haverford BIOL301 Superlab course, who participated in flower trait measurement, genotyping, as well as pollinator observation. Also, we would like to thank Helen White for the help with flower scent analysis.

## Author contributions

FP conceived and designed the project, performed data analysis, and wrote the manuscript. XS, CvV, JC, ML, and WN collected data and performed data analysis. KJ helped with study design, data collection, and manuscript writing. SAK helped with study design, data collection, data analysis, and manuscript writing.

---

**Figure S1.**
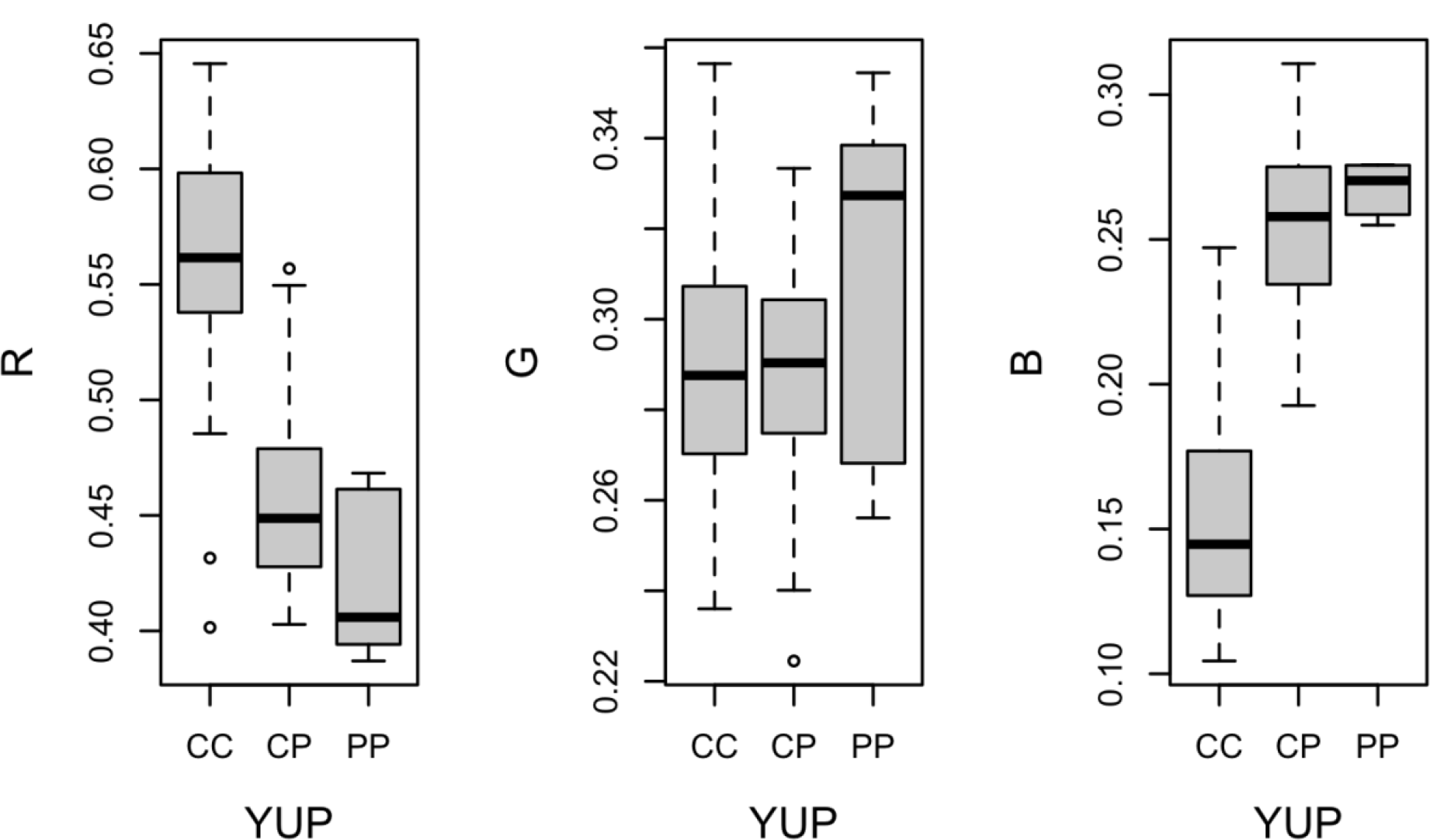
The impact of F2 *YUP* genotype on petal colors. CC: homozygous *M. cardinalis* genotype; PP: homozygous *M. parishii* genotype; PC : heterozygous.

**Figure S2.**
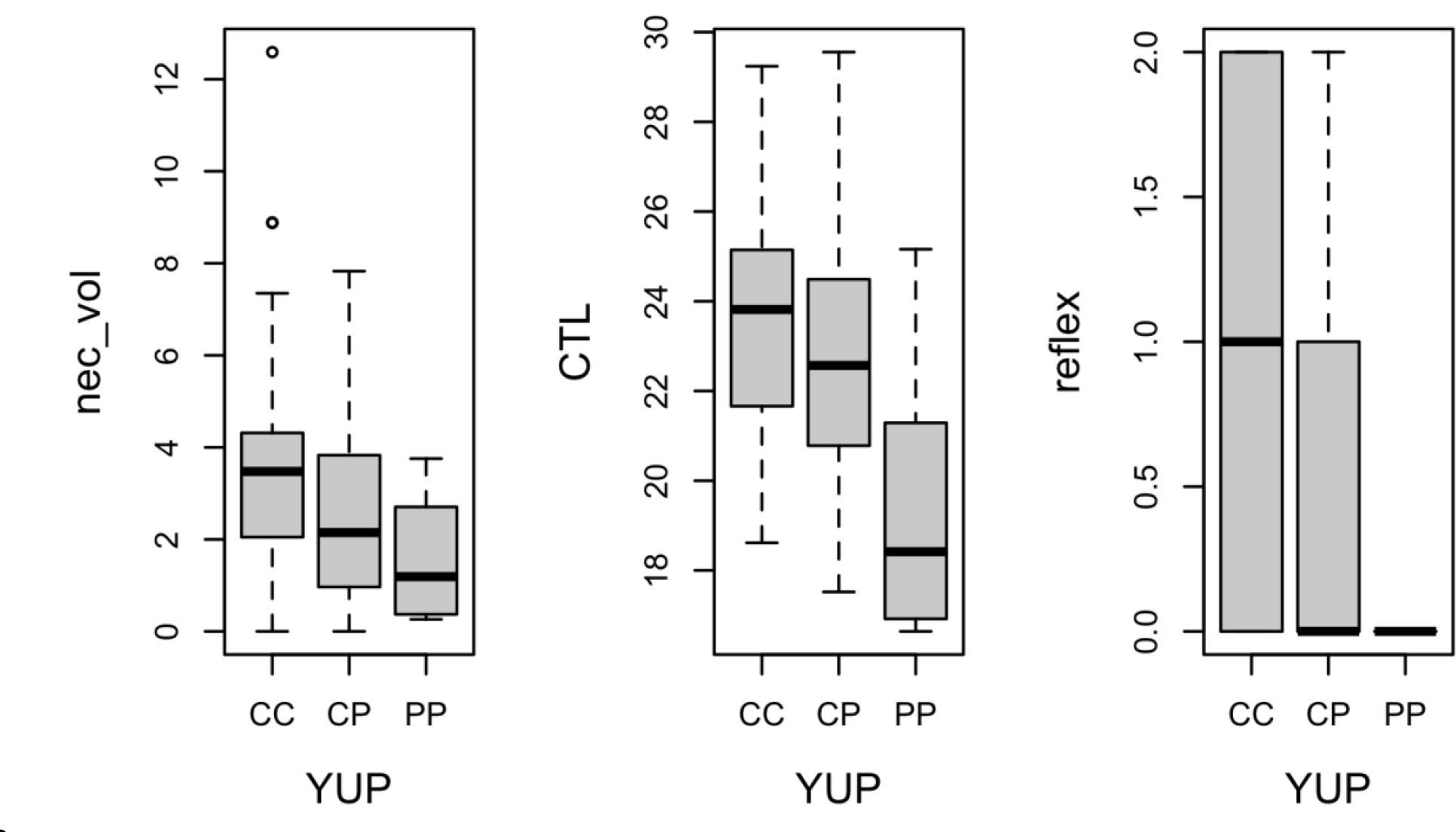
The impact of F2 *YUP* genotype on nectar volume, corolla tube length, and peal reflex. CC: homozygous *M. cardinalis* genotype; PP: homozygous *M. parishii* genotype; PC : heterozygous.

**Figure S3.**
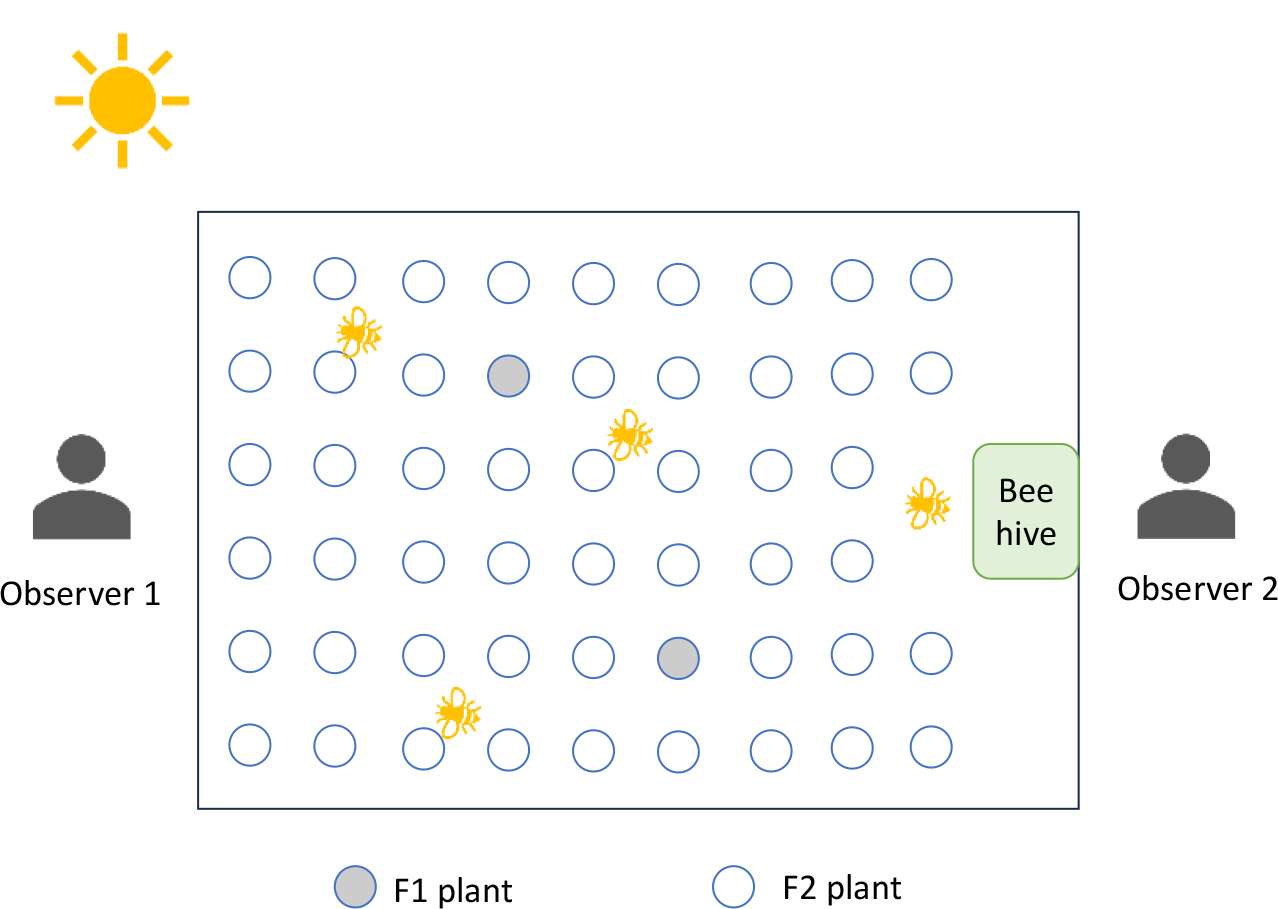
The aerial view of flight cage setup in Pollination Experiment 2.

